# ProCanBio: a database of manually curated biomarkers for Prostate Cancer

**DOI:** 10.1101/2021.06.06.447247

**Authors:** Dikscha Sapra, Harpreet Kaur, Anjali Dhall, Gajendra P. S. Raghava

## Abstract

**Background:** Prostate Cancer is the second lethal malignancy in men worldwide. In the past, numerous research groups investigated the omics profiles of patients and scrutinized biomarkers for the diagnosis and prognosis of prostate cancer. However, information related to the biomarkers is widely scattered across numerous resources in complex textual format, which poses hindrance to understand the tumorigenesis of this malignancy and scrutinization of robust signature. To the best of authors’ knowledge, there is no resource that can consolidate the information contained in all the published literature.

**Results:** Here, we present ProCanBio, a manually curated database that maintains detailed data on 2053 entries of potential prostate cancer biomarkers obtained from 412 publications in user friendly tabular format. Among them, 766 protein-based, 488 RNA-based, 157 genomic mutations, 261 miRNA-based, and 122 are metabolites-based biomarkers. To explore the information in the resource, a web-based interactive platform was developed with searching, and browsing facilities. ProCanBio is freely available and is compatible with most web browsers and devices. Eventually, we anticipated this resource will be highly useful for the research community involved in the area of prostate malignancy.

**Availability:** ProCanBio is available at: https://webs.iiitd.edu.in/raghava/procanbio/

## Introduction

According to the International Agency for research on Cancer, Prostate Cancer is the second most prevalent type of cancer (after Lung cancer) in men, accounting for about 13.5% of the total cancer in 2018 [1]. Prostate Cancer accounts for 1.4 million cases all over world and around 375,000 deaths (6.3% of deaths in males) in 2018 alone [2]. It is the sixth leading cause of cancer related death worldwide [2, 3]. In most cases, Prostate Cancer occurs at the older age, i.e. 65 and above [4].The aggressiveness of the disease is measured by Gleason Score [5]. Gleason Score was introduced in 1966 by Donald F. Gleason. Gleason Score (GS) is a score between 2-10 calculated on the primary and the secondary core of the tumor. The higher the score, the more aggressive is the cancer [6]. Some other non-cancerous prostate conditions include Benign Prostatic Hyperplasia which is one of the most common prostate related diseases, is associated with low urinary tract syndrome. It is not a life threatening disease, but in extreme cases can lead to renal failure. Prostatic intraepithelial neoplasia (PIN) is known to be the most common precursor to prostate cancer. It is further divided into high grade (HGPIN) and low Grade (LGPIN) disease. HGPIN patients are said to be at high risk for prostate cancer [7].

The most common test for the diagnosis and prognosis of Prostate cancer is measurement of Prostate Specific Antigen (PSA) [8]. PSA is present both in normal as well as malignant prostate tissue, however the range of PSA differs in both cases [9]. PSA is very sensitive in detecting PCa but lacks specificity. Elevated levels of PSA are also associated with benign prostate conditions like Benign Prostatic Hyperplasia (BPH), Prostatic Intraepithelial Neoplasia (PIN) and prostatitis which lead to poor specificity [10]. Hence more complex forms of PSA like tPSA (total PSA), fPSA (free PSA), %fPSA have been used for diagnosis. Although these markers enhance diagnosis and prognosis capabilities to a certain extent, however, there is still a need to identify robust biomarkers and drug targets to further improve the prognosis of prostate cancer patients. The need for new biomarkers was also driven by the need to differentiate prostate cancer from BPH, PIN, and prostatitis. The most common treatment for prostate cancer is Androgen Deprivation Therapy (ADT) [11], especially in cases of recurrent and advanced prostate cancer [12]. Over the years there have been a number of studies performed to understand the pathogenesis of this malignancy and to elucidate signatures for prostate cancer [13-16]. This enormous information generated related to prostate cancer biomarkers including genomics, epigenomics, proteomics, metabolomics and peptidomics, etc [17]. But even with the increase in the biomarkers in literature, it is extremely difficult to analyse this information since it is available in unstructured textual format across diverse platforms. This information is widely scattered and there is no repository to store all this information together in a user friendly manner. The absence of structured data format calls the need for ProCanBio.

In the past various databases and prediction tools have been developed for maintaining biomarkers for different types of malignancies such as cervical cancer, liver cancer, breast, colorectal cancer, skin cancer etc [18-33]. To the best of author knowledge, there is no such dedicated database available for maintaining the signatures or biomarkers for prostate cancer. Although, Cancer Proteomics Database includes information related to biomarkers for Prostate cancer, but it only includes Proteomics data and was last updated in 2013 [34]. The Early Detection Research Network provides a list of biomarkers for different type of cancers (including Prostate Cancer), but does not give detailed information about the biomarker, and is not annotated according to individual literature publications.

To fill the gap from the previous studies, in the current study, we developed ProCanBio (https://webs.iiitd.edu.in/raghava/procanbio/) that provides manually curated detailed information from published research articles that related to various biomarkers of prostate cancer. This is a freely accessible database to help researches analyse biomarkers for further use by the scientific community.

## Materials and Methods

### Data Collection

To find all relevant literature related to Prostate Cancer Biomarkers, two keyword searches were performed on PubMed: “(prostate cancer[Title/Abstract]) AND biomarkers[Title/Abstract])” and “(prostate cancer[Title/Abstract]) AND signatures[Title/Abstract])” which yielded 3623 and 422 relevant publications respectively (as analysed on 10th May 2019). From a total of 4045 publications, 112 were common from both the keyword searches and they were removed. A total of 3933 publications were then further analysed to extract information from. After carefully reviewing all articles, reviews, publications which are not available in English language and irrelevant to desired topic were excluded from the study. A total of 412 were left, which were used to extract information about biomarkers.

Extensive information about each biomarker has been found from each publication including PubMed ID, Technical Name, Biomarker Name, Biomarker Basis, Biomolecule, Source, Subjects, Regulation status in Cancerous Conditions, Effect On Pathway, (Hazard Ratio (HR), Odds Ratio (OR), Relative Risk (RR)), experimental conditions, Type of Biomarker, cohort used in the study, performance metrics like Sensitivity, Specificity, Accuracy, ROC-AUC (Receiver Operating Characteristics-Area under the curve), Level of significant (p value), Degree of validity, Clinical Trials and the methods used for analysis were extracted.. Pathway information associated with the obtained from the Enrichr [35].

Each biomarker entry is linked to its original PubMed article from where the biomarker was taken. The Biomarker ID is linked to the GeneCards (version 4.14) [36]. If the biomarker has been assessed in a clinical trial and is registered with the ClinicalTrials (https://clinicaltrials.gov/), it is linked with its NCT number or any other clinical trial.

### Web Interface

ProCanBio developed using APACHE HTTP server which is freely available. The backend is maintained using MySQL (v8) as RDBMS. Front end is developed using PHP(v7), HTML(v5), Javascript (v1.8) and CSS (v3). The server was developed on an Linux machine.

## Results

### Database Architecture

The architecture and overall organisation for ProCanBio is as described in Figure 1

**Figure 1:**
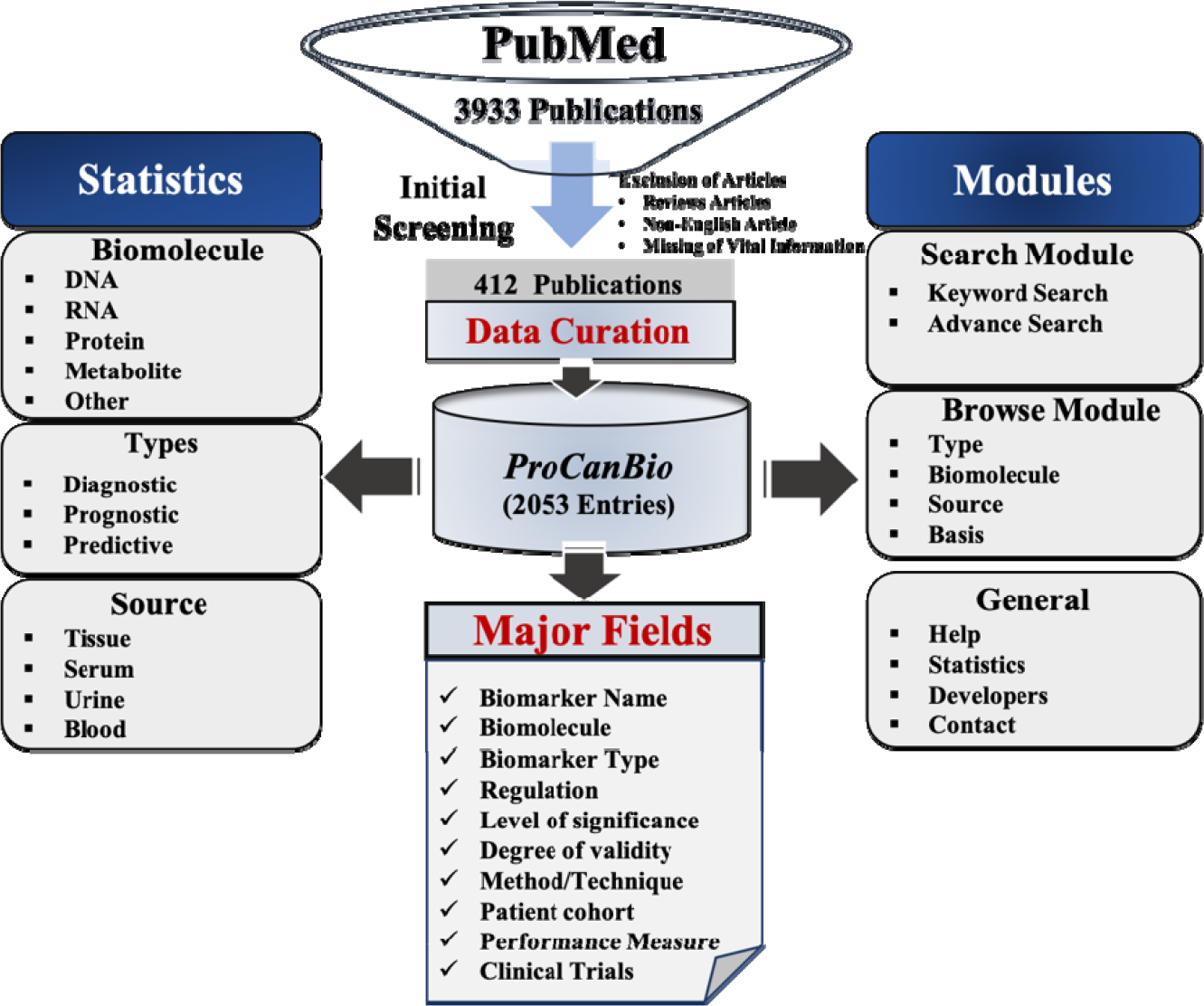
Architecture of ProCanBio

### Biomarkers from the Literature

There are a total of 2053 entries related to biomarkers extracted from 412 publications. Out of which, 1497 are unique biomarkers. The description of each field/column is as follows. “PubMed ID” linked to the original publication from which the information of biomarker was extracted. “Year” represents the yearduring which the study was published; “Biomarker” tells the name of the biomarker/or a group of biomarker that is studied, “Biomarker Basis” gives how the Biomarker was measured - Expression, Concentration, Methylation or Mutation. “Technical Name” refers to the actual name of the Biomarker derived from GeneCard.org. “Biomolecule” provides the bimolecular basis of the biomarker - RNA, DNA, Protein, miRNA, Metabolite etc, “Source” tells from which part of the body this biomarker was extracted - cell lines, tissue, blood, plasma, serum, urine, bone marrow, or semen. “Regulation status in Cancerous Conditions” gives information whether a particular biomarker was observed to be up-regulated or down-regulated in prostate cancer (or the mentioned experimental conditions) along with fold change (difference in the two experimental conditions, if given in original study), “Subjects” tells whether the experiment was performed on Humans, mice or rats. “Odds Ratio/ Hazard Ratio/Relative Risk” tell the Odd Ratio (OR), Hazard Risk (HR) or Relative Ratio (RR) between the two mentioned experimental conditions, “Effect on Pathways” provides information about the different pathways in which a the given biomarker is involved. The pathways information extracted from the Enrichr (Top 5 pathways sorted by adjusted p-values) [35] or GeneCard (pathways with the highest Jaccard Index) or associated publication [36]. “Experiment” refers to the different conditions under which the biomarker was evaluated, “Type of Biomarker” indicates the utility ofgiven biomarker, i.e., diagnostic, prognostic or predictive. Diagnostic biomarkers handle cases where biomarkers are used to segregate prostate cancer from healthy controls, BPH, PIN, and Prostatitis. Prognostic biomarkers are used to identify biomarkers that distinguish Prostate cancer stage, Gleason Score, Metastasis, Biochemical Recurrence, Overall and Disease specific survival. Predictive Biomarkers include Biomarkers which were give information about the effects of therapeutic interventions, particularly, biomarkers that were differentially expressed after a certain therapy. Most common therapies from the database include Docetaxel therapy. “Cohort” gives a description of the patient cohort (population) that was chosen for the study. “sen”, “spec”, “AUC”, “accuracy” refer to the sensitivity, specificity, ROC-AUC, accuracy performance of the biomarkers., respectively in the given experimental conditions. Level of Significance tells the p-value of the experiment. p≤0.05 was considered as significant. “Method Used” provides us the information regarding techniques that were used to perform the experiment in the reported study for biomarker discovery or evaluation likeImmunohistochemistry, qPCR, RT-PCR, fluorescence in situ hybridization (FISH), mass spectrometry, ELISA, Western Blot, MALDI-TOF etc. “Clinical” informs whether the study was a part of a clinical trial, and “Clinical Trial Number” gives the clinical trial ID for the same – either through NCT (National Clinical Trial) or other global trial registrations. “Remarks” include any additional remarks that are to be made about the biomarker card. “Degree of Validity” indicates whether the mentioned biomarker was validated on human patient cohort (in case experiment is performed on cell lines) and it also provides information whether it was validated on an independent dataset or not in the associated study.

In case Biomarker name consists of multiple genes or proteins or miRNAs, the biomarker is either separated by “,”, “;” or “+”. Unless stated otherwise, “,” or “;” indicates that the biomarkers were evaluated individually and more details about each of them are given in the Card. If the biomarkers are separated by “+” that indicates that the biomarker is consists of multiple entities (genes/proteins/miRNAs, etc) and works as a single signature.

In case Biomarker name consists of multiple names, the biomarker is either separated by “,”, “;” or “+”. Unless stated otherwise, “,” or “;” indicates that the biomarkers were evaluated individually and more details about each of them are given in the Card. If the biomarkers are separated by “+” that indicates that the biomarker works together as a signature.

#### Statistics

In the database, stratifying on the basis of the expression of Biomarker-1286, 157 biomarkers were Mutation Based, 218 were Methylation based and 378 were Concentration Based (Figure 2A). Based on the Source, 1069 were extracted from tissues, 354 from Serum, 245 from Blood, 170 from urine, 137 from plasma, 51 from cell lines, 2 from Semen, and 1 from Bone Marrow (Figure 2B). Further, based on the experimental subjects, 2034 Biomarkers were extracted from Humans, 13 from Mice and 5 from Rats. On the basis of Biomolecules: 766 were protein biomarkers, 19 were DNA markers, 488 were RNAs, 261 miRNAs and 122 were Metabolites (Figure 4D). Out of the total biomarker studies, 327 used RT-PCR, 220 used Immunohistochemistry, 97 used ELISA (enzyme-linked immunosorbent assay). The most common biomarker was PCA3 which have 24 entries in the database; Ki-67 appeared 12 times as represented in Table 1.

**Figure 2:**
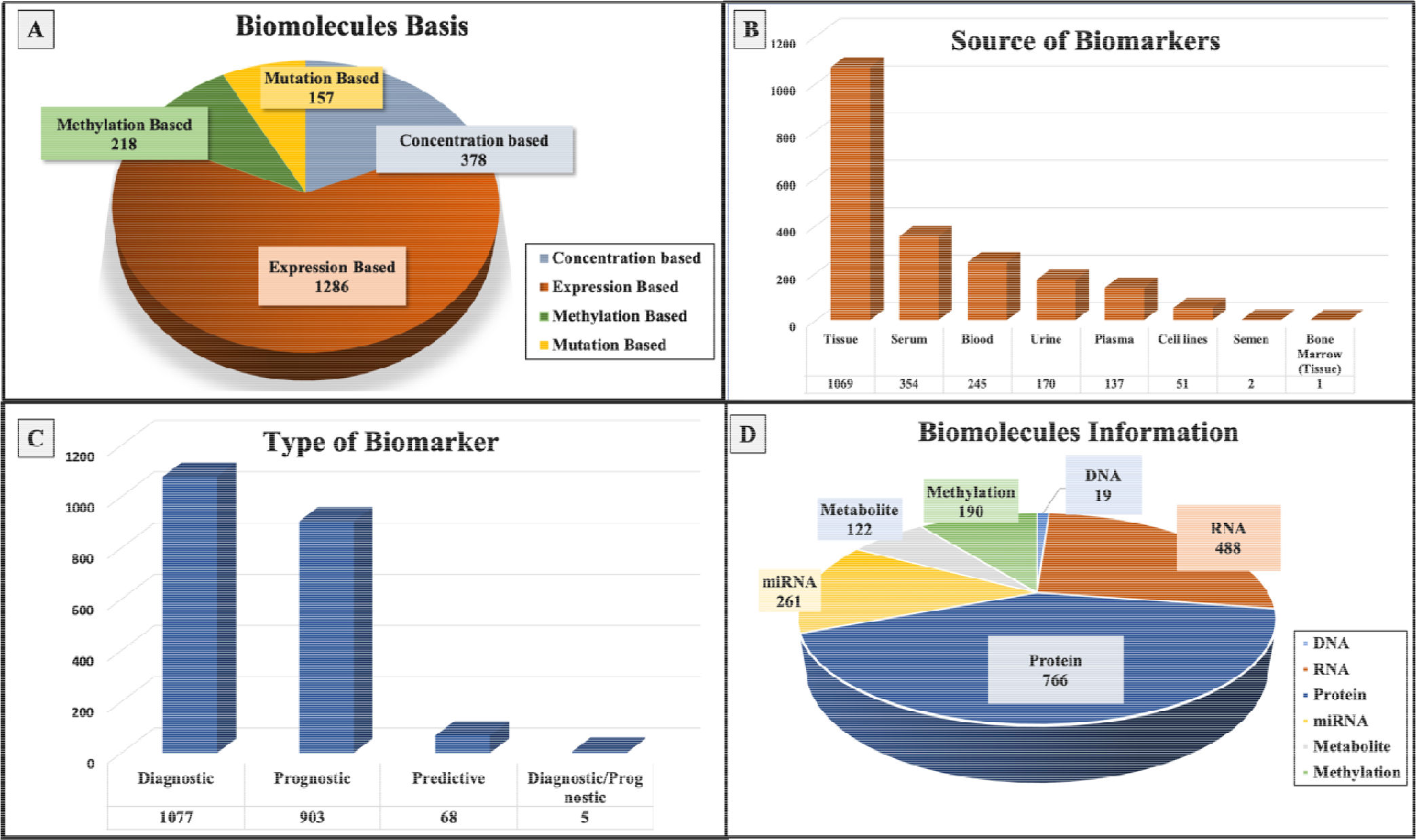
Statistical representation of entries in the PanCanBio: (A) Biomolecule basis, (B) Source of biomarker, (C) Type of Biomarker and (D) Biomolecules Information.

**Figure 3:**
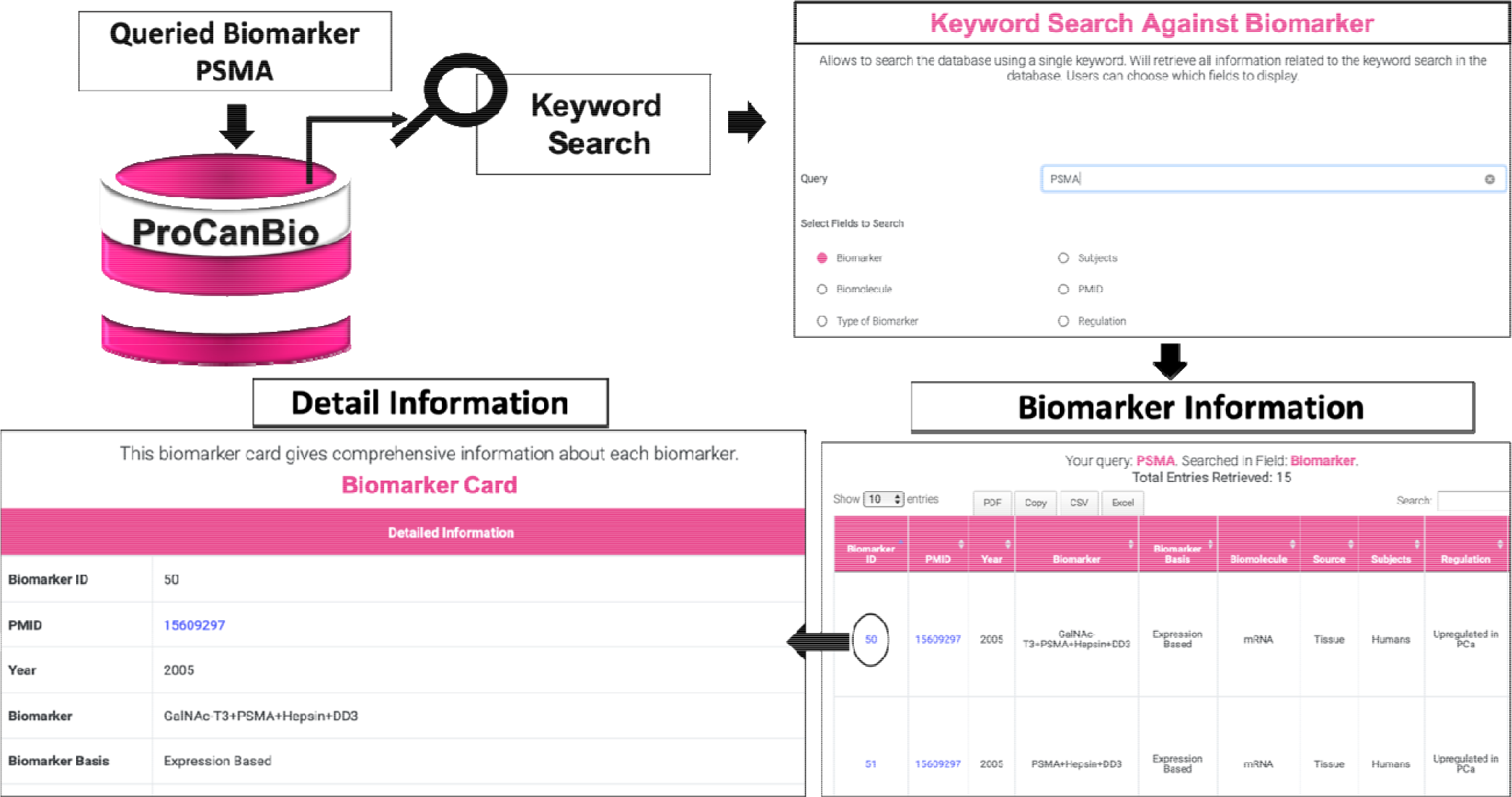
Demonstration of querying and retrieval of information from ProCanBio using keyword search module.

**Figure 4:**
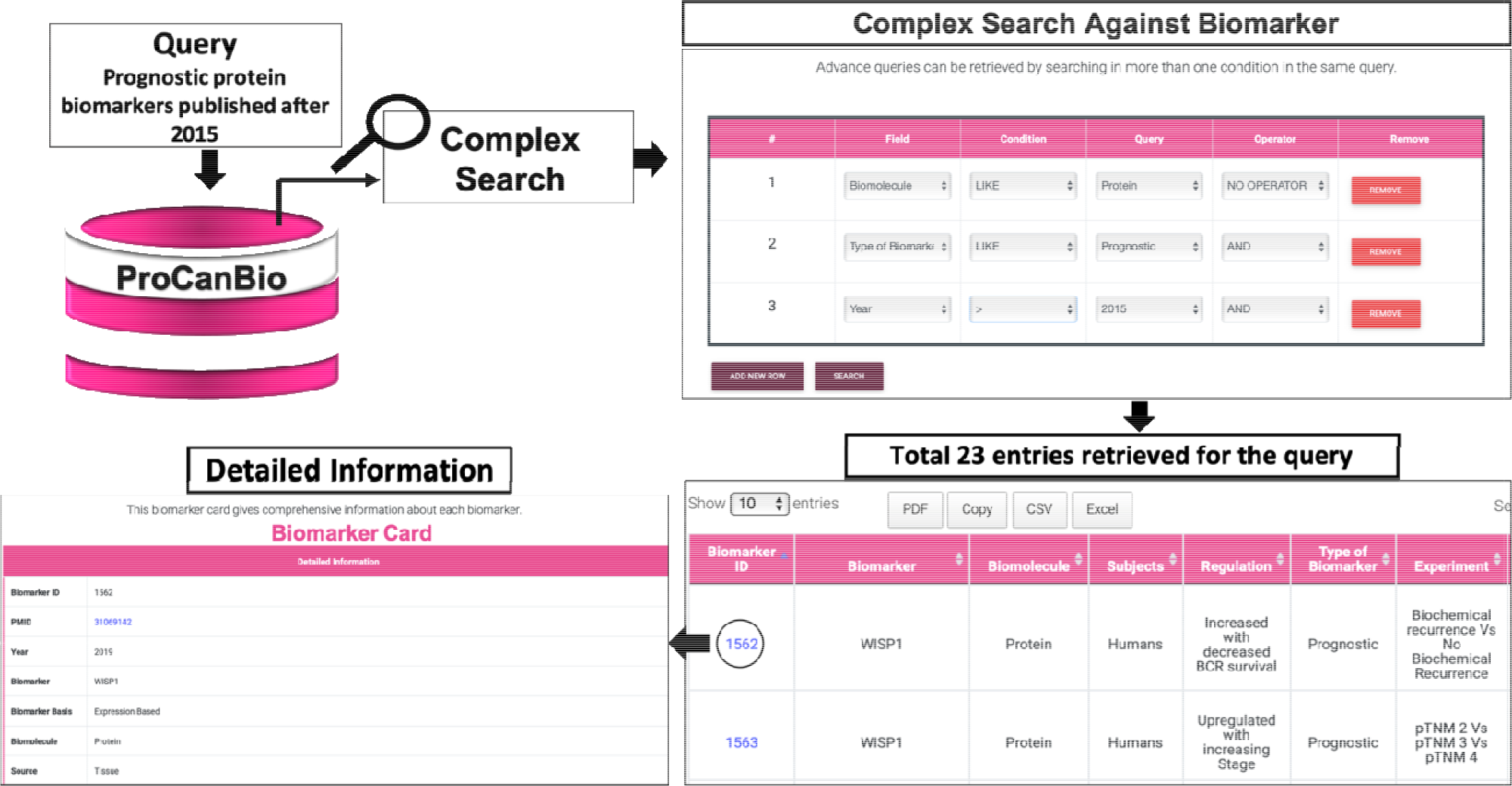
Complex Query Search For Prognostic protein biomarkers published after 2015 in ProCanBio.

**Table 1:** List of top 10 biomarker/signature of prostate cancer with the information such as type of biomolecule and biomarker, number of entries and studies related to the Biomarker.

| Biomarker | Biomolecule | No. of Entries |  |  | No. of Studies |
| --- | --- | --- | --- | --- | --- |
|  |  | Diagnostic | Prognostic | Predictive |  |
| PCA3 | mRNA/Protein | 22 | 2 | 0 | 17 |
| Ki-67 | Protein | 4 | 7 | 1 | 9 |
| Prostate Health Index (phi) | mRNA/Protein | 11 | 0 | 0 | 9 |
| %p2PSA | Protein | 3 | 6 | 0 | 5 |
| miR-141 | miRNA | 3 | 5 | 0 | 8 |
| Prostate Specific Antigen (PSA) | mRNA/Protein | 4 | 1 | 2 | 5 |
| PTEN | mRNA/Protein/DNA | 0 | 7 | 0 | 4 |
| miR-205 | miRNA | 3 | 4 | 0 | 3 |
| Vascular Endothelial Growth Factor (VEGF) | Protein | 2 | 4 | 1 | 5 |
| Methylation status of GSTP1 | mRNA/Methylation | 1 | 6 | 0 | 5 |

### Querying the Database

The ProCanBio has the facilities to retrieve the data using many different search options like Searching and Browsing: including Keyword Search, Complex Search, Type of Biomarker, Type of Biomolecule, Source, Basis of Biomarker

A. **Search Tools:** These tools take an input from user and perform search queries on the database which are in agreement to the keyword(s) entered by the user.
  1. Keyword Search: It allows to search the database using a single keyword. It will retrieve all information related to the keyword search in the database. Fields can be selected according to name of Biomarker, Biomolecule, Type of Biomarker, Subjects, PMID and Regulation conditions, etc. The search is not case sensitive, so as it takes queries in all cases of the user. Users can choose which fields to display by ticking the provided check boxes.
  2. Complex Search: It allows user to access the database when ProCanBio query with more than one keyword. The complex query can be searched on Biomolecules, Source of Biomarker, Type of Biomarkers, Subjects and Year of Publication. For Numeric type operator (Year), field option can be set to =, <, > to retrieve queries. The rest of the operators (excluding year) are string type operators and can be queried using “like” field option. Condition field contains all permissible values for the operator field. Thus, rows can be added and deleted to be more complex queries. The Query field allows to join different individual queries by “and”, “not” or “or” operations. For example, if one has to search all entries from database such that we have to find proteins with prognostic capabilities for Prostate Cancer, we can do so by entering “Biomolecule” in the first row operator, with LIKE field, and “protein” condition then add new row, and select “Type of Biomolecule” operator, with “Like” field and “prognostic” in the condition field along with “and” in the Query field (since we want our result to satisfy both these conditions).
B. **Browse Tools:** These tools help to query the database on certain main keywords in an orderly manner. The user can query using four main tools - (i) Type of Biomarker (ii) Biomolecule (iii) Source (iv) Biomarker Basis. On clicking on all these options, user is redirected to a page which will provide different categories for all above mentioned four tools. User can selected any one of them to see all related entries for the keyword in the database.

While viewing query results for both Search and Browse, user can select how many entries to be shown on one page, and allows search within the queried dataset. When a keyword is entered, if any of the rows contain that keyword in the row, then that row would be retained. The results can be downloaded in the form of excel (.xls), comma separated values (.csv), or a PDF file. To view the detailed information regarding the biomarker, ID column should be selected which provides a Biomarker Card which contains all the information regarding that particular entry in the database.

In the general tab, user can view Help dropdown which has detailed instructions on how to use the server. The statistics dropdown gives statistics about the various biomarkers in ProCanBio. Developers and Contact dropdown shows information about the developers of ProCanBio.

## Working of PanCanBio Database

### Case Study I: PSMA as Biomarker

ProCanBio can facilitate the user for easy query and browse the database without having to go through hundreds of entries to find the relevant information. Here we illustrate the results for query against Prostate Specific Membrane Antigen (PSMA). After performing keyword search against ‘PSMA’, we find there are 15 unique entries in ProCanBio from 6 research publications as shown in Figure 3. This information tells us it has been used for diagnostic purposes (mostly against benign prostatic Hyperplasia). It has been extracted from a variety of samples including tissue, serum and urine. It has been validated only once on independent datasets. We can observe that PSMA mean expression is higher in Prostate Cancer as compared to controls. Upon clicking the Biomarker ID, we can see detailed information for PSMA including Odds Ratio, Performance, Special Remarks and Degree of Validity and Clinical Trial Number (none in this case, since PSMA is not studied in any clinical trials).

### Case Study II: Published Prognostic Protein Biomarkers after 2015

To perform complex queries, we show a case study to show all prognostic protein biomarkers published after the year 2015. By performing this query, we retrieve 23 unique entries. Out of these 23 entries, 5 focus on biochemical recurrence and 7 focus on overall survival. These are taken from either tissues or Serum of patients. Some of the important biomarkers that are retrieved include Prohibitin, WISP1, Ki-67, GLUT1 and AZGP1. The query performed to find these results is as shown in Figure 4. Similar to these case studies, more queries can be performed to answer similar questions related to Prostate Cancer Biomarkers.

## Discussion

ProCanBio provides cumulative information about biomarkers related to Prostate Cancer. The information from each paper is manually curated before adding it into the database. Extensive information was collected from each published article to provide detailed information to the user - providing not only the regulation status and fold change, but more information like patient cohort, degree of validity, methods used to analyse the biomarker, performance metrics etc. Besides, this database is extremely exhaustive since it focuses on genomics, proteomics and metabolomics instead of focusing on one particular aspect like Cancer Proteomics Database [34]. ProCanBio encompass a total 2053 entries, with 1497 unique biomarkers. Tabular display of information and user friendly interface makes it easy for people from all backgrounds to use this resource. It is freely available to the scientific community without having to login to any platform. The interface is extremely user friendly and one can easily browse and search through the biomarkers database.

### Comparison with other Resources

Currently, there are not many resources available for biomarkers related to Prostate Cancer. One such resource is Cancer Proteomics Database. Although, it contains proteomics biomarkers, but, it lacks genomic, metabolomic and epigenomic, which were provided by the ProCanBio in addition to proteins based biomarkers. Further, Cancer Proteomics Database extracted information from 143 articles upto the year 2013, whereas ProCanBio covers 412 articles. Though Cancer Proteomics Database does tell some information about the biomarkers, it lacks vital information like the performance of the biomarker, significant level, degree of validity, etc.

The another resource, Early Detection Research Network (ERDN), National Cancer Institute (NCI) publishes a list of biomarkers for several types of cancer including Lung, Prostate, Liver, Breast, Lung etc. They provide information for each biomarker, its aliases, which cancers are they over expressed in, and if there are any studies related to the biomarker published by their own organisation. But no further details about the biomarkers is given.

ProCanBio can be used to find supporting evidence about a specific biomarker from the published literature to further use it for experimental validation. The motivation to develop ProCanBio was to provide a freely available, comprehensive database regarding all related literature on a disease that affects more than 1.27 million lives every year. We anticipate, this resource will be useful for the scientific community actively involved in the elucidation of biomarkers for the prostate cancer.

### Applications of the Database

This database can be used for a variety of applications that can help the research and scientific community. Some of them are listed as follows:

1. To the best of the author’s knowledge, no other database currently provides information about various signatures and biomarkers for prostate cancer from such as wide fields including genomics, epigenomics and metabolomics.
2. Easy to comprehend User interface means people with minimal knowledge can use this database.
3. Pubmed, GeneCard, ClinicalTrials, Enrichr are all linked to the platform, making it easy for the reader to connect all the different platforms and find the relevant information related to a particular biomarker.
4. ‘Complex Search’ tool allows users to find answers to complex queries which can help narrow down research already done on a particular biomarker.
5. Browsing and Searching tools for better understanding and comprehension of the existing literature.
6. ProCanBio is particularly useful to retrieve detailed supporting evidence from published literature in order to select a particular biomarker for further research on prostate cancer.

### Future Development

Since the prostate cancer is one of major concern worldwide, thus, scientific community is continuously working in this field. With the availability of more articles with sufficient information for prostate cancer in the future, our first aim will be to update the resource with available information. Besides, we will also expand information from biomarkers to the drugs and therapeutic options.

## Acknowledgement

D.S., H.K.,and A.D. are grateful to the Department of Biotechnology (DBT), India, the Council of Scientific and Industrial Research (CSIR), India, and the Department of Science Technology (DST), India, for providing fellowships, respectively.

## Availability of data and material

ProCanBio is available at: https://webs.iiitd.edu.in/raghava/procanbio/

## Author’s Contribution

**Conception and design:** Dikscha Sapra, Harpreet Kaur, and Gajendra P. S. Raghava

**Development of methodology:** Dikscha Sapra, Harpreet Kaur, and Gajendra P. S. Raghava

**Acquisition of data:** Dikscha Sapra

**Analysis and interpretation of data and results:** Dikscha Sapra, Harpreet Kaur, Anjali Dhall, Gajendra P. S. Raghava

**Webserver Development:** Dikscha Sapra, Harpreet Kaur, Anjali Dhall

**Writing, reviewing, and revision of the manuscript:** Dikscha Sapra, Harpreet Kaur, Anjali Dhall, Gajendra P. S. Raghava

**Supervision and Coordination of the Project:** Gajendra P. S. Raghava

## References

1. Rawla P. Epidemiology of Prostate Cancer, World J Oncol 2019;10:63–89.

2. Sung H, Ferlay J, Siegel RL et al. Global Cancer Statistics 2020: GLOBOCAN Estimates of Incidence and Mortality Worldwide for 36 Cancers in 185 Countries, CA Cancer J Clin 2021;71:209–249.

3. Jemal A, Siegel R, Xu J et al. Cancer statistics, 2010, CA Cancer J Clin 2010;60:277–300.

4. Droz JP, Balducci L, Bolla M et al. Management of prostate cancer in older men: recommendations of a working group of the International Society of Geriatric Oncology, BJU Int 2010;106:462–469.

5. Verma S, Rajesh A, Morales H et al. Assessment of aggressiveness of prostate cancer: correlation of apparent diffusion coefficient with histologic grade after radical prostatectomy, AJR Am J Roentgenol 2011;196:374–381.

6. Mellinger GT, Gleason D, Bailar J, 3rd. The histology and prognosis of prostatic cancer, J Urol 1967;97:331–337.

7. Brawer MK. Prostatic intraepithelial neoplasia: an overview, Rev Urol 2005;7 Suppl 3:S11–18.

8. Atan A, Guzel O. How should prostate specific antigen be interpreted?, Turk J Urol 2013;39:188–193.

9. Adhyam M, Gupta AK. A Review on the Clinical Utility of PSA in Cancer Prostate, Indian J Surg Oncol 2012;3:120–129.

10. Nadler RB, Humphrey PA, Smith DS et al. Effect of inflammation and benign prostatic hyperplasia on elevated serum prostate specific antigen levels, J Urol 1995;154:407–413.

11. Huggins C, Hodges CV. Studies on prostatic cancer: I. The effect of castration, of estrogen and of androgen injection on serum phosphatases in metastatic carcinoma of the prostate. 1941, J Urol 2002;168:9–12.

12. Moul JW, Kibel AS, Roach M, 3rd et al. Indications and practice with androgen deprivation therapy, Urology 2011;78:S478–481.

13. Pretorius ME, Waehre H, Abeler VM et al. Large scale genomic instability as an additive prognostic marker in early prostate cancer, Cell Oncol 2009;31:251–259.

14. Hessels D, Smit FP, Verhaegh GW et al. Detection of TMPRSS2-ERG fusion transcripts and prostate cancer antigen 3 in urinary sediments may improve diagnosis of prostate cancer, Clin Cancer Res 2007;13:5103–5108.

15. Kosari F, Cheville JC, Ida CM et al. Shared gene expression alterations in prostate cancer and histologically benign prostate from patients with prostate cancer, Am J Pathol 2012;181:34–42.

16. Hutchinson LM, Chang EL, Becker CM et al. Use of thymosin beta15 as a urinary biomarker in human prostate cancer, Prostate 2005;64:116–127.

17. Sallam RM. Proteomics in cancer biomarkers discovery: challenges and applications, Dis Markers 2015;2015:321370.

18. Kaur H, Kumar R, Lathwal A et al. Computational resources for identification of cancer biomarkers from omics data, Brief Funct Genomics 2021.

19. Agarwal SM, Raghav D, Singh H et al. CCDB: a curated database of genes involved in cervix cancer, Nucleic Acids Res 2011;39:D975–979.

20. Kaur H, Bhalla S, Kaur D et al. CancerLivER: a database of liver cancer gene expression resources and biomarkers, Database (Oxford) 2020;2020.

21. Zhang X, Sun XF, Cao Y et al. CBD: a biomarker database for colorectal cancer, Database (Oxford) 2018;2018.

22. Dingerdissen HM, Bastian F, Vijay-Shanker K et al. OncoMX: A Knowledgebase for Exploring Cancer Biomarkers in the Context of Related Cancer and Healthy Data, JCO Clin Cancer Inform 2020;4:210–220.

23. Perez-Granado J, Pinero J, Furlong LI. ResMarkerDB: a database of biomarkers of response to antibody therapy in breast and colorectal cancer, Database (Oxford) 2019;2019.

24. Chu YW, Chien CH, Sung MI et al. dBMHCC: A comprehensive hepatocellular carcinoma (HCC) biomarker database provides a reliable prediction system for novel HCC phosphorylated biomarkers, PLoS One 2020;15:e0234084.

25. Zuo Z, Hu H, Xu Q et al. BBCancer: an expression atlas of blood-based biomarkers in the early diagnosis of cancers, Nucleic Acids Res 2020;48:D789–D796.

26. Nalejska E, Maczynska E, Lewandowska MA. Prognostic and predictive biomarkers: tools in personalized oncology, Mol Diagn Ther 2014;18:273–284.

27. Butti MD, Chanfreau H, Martinez D et al. BioPlat: a software for human cancer biomarker discovery, Bioinformatics 2014;30:1782–1784.

28. Terkelsen T, Krogh A, Papaleo E. CAncer bioMarker Prediction Pipeline (CAMPP)-A standardized framework for the analysis of quantitative biological data, PLoS Comput Biol 2020;16:e1007665.

29. Dhall A, Patiyal S, Kaur H et al. Computing Skin Cutaneous Melanoma Outcome From the HLA-Alleles and Clinical Characteristics, Front Genet 2020;11:221.

30. Bhalla S, Kaur H, Dhall A et al. Prediction and Analysis of Skin Cancer Progression using Genomics Profiles of Patients, Sci Rep 2019;9:15790.

31. Kaur H, Bhalla S, Raghava GPS. Classification of early and late stage liver hepatocellular carcinoma patients from their genomics and epigenomics profiles, PLoS One 2019;14:e0221476.

32. Bhalla S, Chaudhary K, Kumar R et al. Gene expression-based biomarkers for discriminating early and late stage of clear cell renal cancer, Sci Rep 2017;7:44997.

33. Kaur H, Dhall A, Kumar R et al. Identification of Platform-Independent Diagnostic Biomarker Panel for Hepatocellular Carcinoma Using Large-Scale Transcriptomics Data, Front Genet 2019;10:1306.

34. Arntzen MO, Boddie P, Frick R et al. Consolidation of proteomics data in the Cancer Proteomics database, Proteomics 2015;15:3765–3771.

35. Kuleshov MV, Jones MR, Rouillard AD et al. Enrichr: a comprehensive gene set enrichment analysis web server 2016 update, Nucleic Acids Res 2016;44:W90–97.

36. Stelzer G, Rosen N, Plaschkes I et al. The GeneCards Suite: From Gene Data Mining to Disease Genome Sequence Analyses, Curr Protoc Bioinformatics 2016;54:1 30 31-31 30 33.

